# Detection of reproducible liver cancer specific ligand-receptor signaling in blood

**DOI:** 10.1101/2023.09.25.559274

**Authors:** Aram Safrastyan, Damian Wollny

## Abstract

Cell-cell communication mediated by ligand-receptor interactions (LRI) is critical to coordinating diverse biological processes in homeostasis and disease. Lately, our understanding of these processes has greatly expanded through the inference of cellular communication, utilizing RNA extracted from bulk tissue or individual cells. Considering the challenge of obtaining tissue biopsies for these approaches, we considered the potential of studying cell-free RNA obtained from blood. To test the feasibility of this approach, we used the BulkSignalR algorithm across 295 cell-free RNA samples and compared the LRI profiles across multiple cancer types and healthy donors. Interestingly, we detected specific and reproducible LRIs particularly in the blood of liver cancer patients compared to healthy donors. We found an increase in the magnitude of hepatocyte interactions, notably hepatocyte autocrine interactions in liver cancer patients. Additionally, a robust panel of 30 liver cancer-specific LRIs presents a bridge linking liver cancer pathogenesis to discernible blood markers. In summary, our approach shows the plausibility of detecting liver LRIs in blood and builds upon the biological understanding of cell-free transcriptomes.

## Introduction

Cancer remains one of the most pressing healthcare challenges globally, being the second leading cause of death worldwide (GBD 2015 Mortality and Causes of Death Collaborators, 2016). Numerous studies have shown that early cancer detection significantly improves the survival rate, emphasizing the importance of improved detection methods (Crosby et al., 2022). A promising method for minimally invasive yet highly informative diagnostics is liquid biopsy. This methodology focuses on analyzing body fluids - primarily blood - utilizing various omics techniques such as proteomics, genomics, and, notably, transcriptomics (Heitzer et al., 2019). In particular, the analysis of cell-free RNA is increasingly promising (Cabús et al., 2022) due to the fact that transcriptomic signatures can reveal tissue and cell-type specificity which would greatly aid diagnostics (Jin et al., 2023; Koh et al., 2014; Vong et al., 2021; Vorperian et al., 2022; Zaporozhchenko et al., 2018). Beyond diagnostics, cell-free RNA can in principle offer many insights into cellular processes of cells throughout the body since RNAs are constantly being shed into the bloodstream (Ibarra et al., 2020; Zaporozhchenko et al., 2018). Yet, to which extent we can learn about inter- and intracellular processes solely by investigating the limited subset of cell transcriptomes that enter into the bloodstream is currently unknown.

In recent years, numerous approaches have focussed on mapping ligand-receptor interactions as they are crucial for comprehending cellular responses and intercellular communication networks (Cabello-Aguilar et al., 2020; Dimitrov et al., 2022; Efremova et al., 2020; Lu et al., 2022). In particular, single-cell RNA sequencing (scRNA-seq) technology enables the measurement of ligand and receptor expression across various cell types, facilitating the systematic decoding of intracellular communication for the maintenance of homeostasis but also in cancerogenesis (Ghoshdastider et al., 2021; Ramilowski et al., 2015; Zhou et al., 2017). In order to experimentally gain these insights, however, tissue biopsies are needed which are difficult to extract and only provide a snapshot in time. Thus, we asked the question as to which degree one could observe LRI differences between cancer and normal tissue solely by exploring the cell-free RNA found in the blood of patients and healthy donors.

In this proof-of-concept study, using the BulkSignalR algorithm (Villemin et al., 2023), we queried LRIs in close to 300 blood samples, including those from cancer patients and healthy donors. In our analysis of all the samples, liver cancer samples notably distinguished themselves. We showed not only the possibility of inferring relevant LRIs from cell-free transcriptomes in blood from liver cancer patients but also highlighted an increase in the number of hepatocyte interactions associated with hepatocytes in these patients. Furthermore, we curated a panel of 30 highly robust LRIs in the cell-free transcriptome specific to liver cancer. Within this panel, we find previously documented liver- and liver cancer-relevant marker gene ligands SERPINC1 and GPC3 LRIs to be specific and unique to liver cancer blood samples and thus serve as potentially potent biomarkers.

## Materials and Methods

### Detection of LRIs

We used publicly available RNA-seq datasets generated by Chen *et al*. (Chen et al., 2022) and Zhu *et al*. (Zhu et al., 2021) to study the LRIs in blood cell-free transcriptomes. The count matrices of raw reads were downloaded from Gene Expression Omnibus (GEO) with the ascension numbers GSE174302 and GSE142987. In total, 295 blood samples from five types of solid tumors and healthy donors were analyzed (Table 1). Liver cancer (LC) blood samples (n=62) were mainly drawn from hepatocellular carcinoma (HCC) patients, with only eight samples drawn from intrahepatic cholangiocarcinoma (ICC) patients (Chen et al., 2022; Zhu et al., 2021). Additionally, more than 60% of liver cancer patients had chronic hepatitis B (CHB) infection (Chen et al., 2022).

**Table 1:** Main characteristics of the RNA-seq datasets used in the study. HD, healthy donor; LC, liver cancer; STAD, stomach adenocarcinoma; LUAD, lung adenocarcinoma; CRC, colorectal cancer; ESCA, esophageal cancer.

| Dataset | HD | LC | STAD | LUAD | CRC | ESCA | Accession number | Reference |
| --- | --- | --- | --- | --- | --- | --- | --- | --- |
| Chen *et al.* (2022) | 46 | 27 | 37 | 35 | 54 | 31 | GSE174302 | (Chen et al., 2022) |
| Zhu *et al.* (2021) | 30 | 35 | NA | NA | NA | NA | GSE142987 | (Zhu et al., 2021) |
| Single-cell RNA-seq | NA | 17,392 cells/5 patients | NA | NA | NA | NA | GSE149614 | (Lu et al., 2022) |
| Bulk RNA-seq | NA | 35 | NA | NA | NA | NA | GSE124535 | (Jiang et al., 2019; Zeng et al., 2020) |

In order to infer LRIs, the count matrices were separately (per biological condition) prepared for subsequent analysis using the function “prepareDataset” from the R (version 4.1.2) (R Core Team, 2021) package BulkSignalR (version 0.0.9) (Villemin et al., 2023). Next, the function “learnParameters” from the package BulkSignalR was employed to estimate the statistical model parameters and finally, with the function “initialInference” from the package BulkSignalR LRIs are inferred and stored in a BSRInference object. Dataframes of inferred LRIs were extracted from the BSRInference objects using the function “LRInter” from the package BulkSignalR. A threshold of 0.1% FDR was applied, as described in the original study (Villemin et al., 2023).

A count matrix of bulk RNA-seq dataset of CHB HCC tissue samples (Jiang et al., 2019; Zeng et al., 2020) was downloaded from GEO under the ascension number GSE124535. The count matrix contained FPKM values of 35 HCC liver tissue samples and was used to infer LRIs with the R package BulkSignalR as described with the parameter “normalize=FALSE” in the “prepareDataset” function. In order to get only high-confidence LRIs, a threshold of 0.1% FDR and a minimum ligand-receptor correlation value of 0.5 were applied.

Finally, a single-cell RNA-seq dataset of HCC patient liver tissue samples (Lu et al., 2022) was downloaded from the GEO database under the ascension number GSE149614. After selecting cells extracted from primary tumor sites of CHB patients, there remained 17,392 cells from five patients. To annotate the cells, we used the R Bioconductor (Huber et al., 2015) package SingleR (version 1.8.0) (Aran et al., 2019) and a healthy liver single-cell RNA-seq dataset (MacParland et al., 2018) as reference. The latter was downloaded from the GEO database and included log2CPM values of 8,444 cells from five healthy donor liver tissues. After collapsing the annotations for hepatocytes, macrophages, T cells and liver sinusoidal endothelial cells (LSECs), 11 cell-type annotations remained and, together with the liver cancer scRNA-seq data were used as input for the “SingleR” function of the SingleR package with the parameter “de.method=“wilcox”“. Then, the annotated liver cancer scRNA-seq data was normalized with the function “NormalizeData” of the Seurat R package (version 4.3.0.1) (Hao et al., 2021). Finally, LRIs were inferred from the annotated and normalized HCC single-cell RNA-seq data using the function “liana_wrap” from the LIANA R package (version 0.1.12) (Dimitrov et al., 2022) with the parameter “resource = “LRdb”“as the package BulkSignalR also uses the database LR*db* (Cabello-Aguilar et al., 2020). In order to acquire only high-confidence LRIs, we applied a threshold of 0.05 for the CellPhoneDB *p*-values (Dimitrov et al., 2022; Efremova et al., 2020) and a threshold of 0.5 for the correlations between ligands and receptors (Dimitrov et al., 2022; Villemin et al., 2023).

The LRIs inferred from Chen *et al*. liquid biopsy datasets were used as input for the “UpsetR” function from the UpsetR R package (version 1.4.0) (Conway et al., 2017) to generate UpSet plots. Only the first ten intersections ordered by size were shown for visualization purposes. The intersection between liver cancer samples from Chen *et al*. and Zhu *et al*. datasets was visualized using the “venn” function from the R package ggvenn (version 0.1.10) (Yan, 2023) with the parameter “auto_scale=T”.

The total number of LRIs identified in Chen *et al*. and Zhu *et al*. datasets was visualized using the R package ggplot2 (version 3.4.3) (Hadley, 2016) and ggpubr (version 0.6.0) (Kassambara, 2023). The relationship between tissue bulk and single-cell RNA-seq LRIs was visualized with the “ggvenn” function from the ggvenn R package with the parameter “auto_scale=TRUE”.

### Inference of cellular interactions

In order to associate the identified LRIs in the cell-free transcriptome with source and target cell types, we used the scRNA-seq results where each identified LRI was assigned to a source and target cell type found in the liver. First, the healthy donor blood LRIs were removed from the lists of LRIs identified in the blood of cancer patients. Then, the scRNA-seq LRI dataframe was filtered by LRIs found in each cell-free data type. Then, in each resulting dataframe the number of interactions between cell types was calculated and the top ten interactions were selected. Finally, the selected cell interactions were used to construct cellular networks with the R package circlize (version 0.4.14) (Gu et al., 2014) using the function “chordDiagramFromMatrix”. Hepatocyte autocrine interactions were highlighted where present with a dashed line.

### Generation of an LC-specific and reproducible panel of LRIs

To generate a panel of LC-specific and reproducible LRIs, we first excluded every LRI found in liver cancer cell-free dataset of Chen *et al*. that could also be found in Chen *et al*. healthy donor, esophageal cancer, stomach adenocarcinoma, colorectal cancer and lung adenocarcinoma cell-free datasets. Then, we intersected the remaining LRIs with those found in Zhu *et al*. liver cancer cell-free dataset and finally excluded LRIs found also in Zhu *et al*. healthy donor cell-free dataset. The final list included 30 LRIs which were used to generate corresponding alluvial diagrams with the function “alluvialplot” from the BulkSignalR package. Considering that LRIs may be associated with multiple biological pathways, we visualized the largest pathways per LRI.

To further contextualize the identified LRIs, the online portal CITE (Crosstalk Interactions within Tumor microenvironment; https://cite.genome.sg/) (Ghoshdastider et al., 2021) was used to acquire the estimated Relative Crosstalk (RC) score for each LRI in HCC. RC scores represent the relative concentration of LR complexes in cancer and stromal cell compartments in the tumor microenvironment and the directionality of interactions between compartments (Ghoshdastider et al., 2021). The portal contained information about 11 of the identified 30 LRIs and after retrieving the RC scores, they were used to generate a heatmap using the function “pheatmap” from the R package pheatmap (version 1.0.12) (Kolde, 2019).

In order to visualize the expression patterns of the ligands SERPINC1 and GPC3 across tumor tissues, the online portal GEPIA2 (http://gepia2.cancer-pku.cn/) was employed. After retrieving the expression values, they were used for visualization with the packages ggplot2 and ggpubr.

## Results and Discussion

### Extensive LRI detection in liver cancer cell-free transcriptome

To evaluate the possibility of detecting ligand-receptor interactions (LRIs) in the cell-free transcriptome in blood, we utilized the BulkSignalR package and tested it using 295 liquid biopsy samples from healthy donors, liver cancer, esophageal cancer, stomach adenocarcinoma, colorectal cancer and lung adenocarcinoma sourced from the publicly available datasets of Chen *et al*. (Chen et al., 2022) and Zhu *et al*. (Zhu et al., 2021). We detected statistically significant LRIs in both datasets, with numbers ranging from 67 to 455 (Suppl. Figure 1A; Suppl. File 1A). Notably, both cell-free liver cancer cohorts displayed a markedly higher number of total and unique LRIs compared to other cell-free RNA samples from other types of cancer (Suppl. Figure 1A; Figure 1A), underscoring the pronounced increase in tissue signal presence in blood during liver cancerogenesis. Most of the LRIs identified in cell-free RNA liver cancer data were reproducible in both cohorts (Figure 1B), reducing the likelihood of them being attributed to random noise. This unique feature of liver cancer results can be explained by the pronounced contribution of the liver to the cell-free transcriptome (Larson et al., 2021) which means that any pathological changes within the liver are prominently reflected in blood samples (Morlion et al., 2023; Vong et al., 2021). Earlier studies have highlighted detectable alterations in the cell-free transcriptome during liver diseases, with particular shifts in cell-type signals (Vorperian et al., 2022). Hence, this finding indicates that liver cancer is a promising use case for studying LRI changes during carcinogenesis.

**Figure 1:**
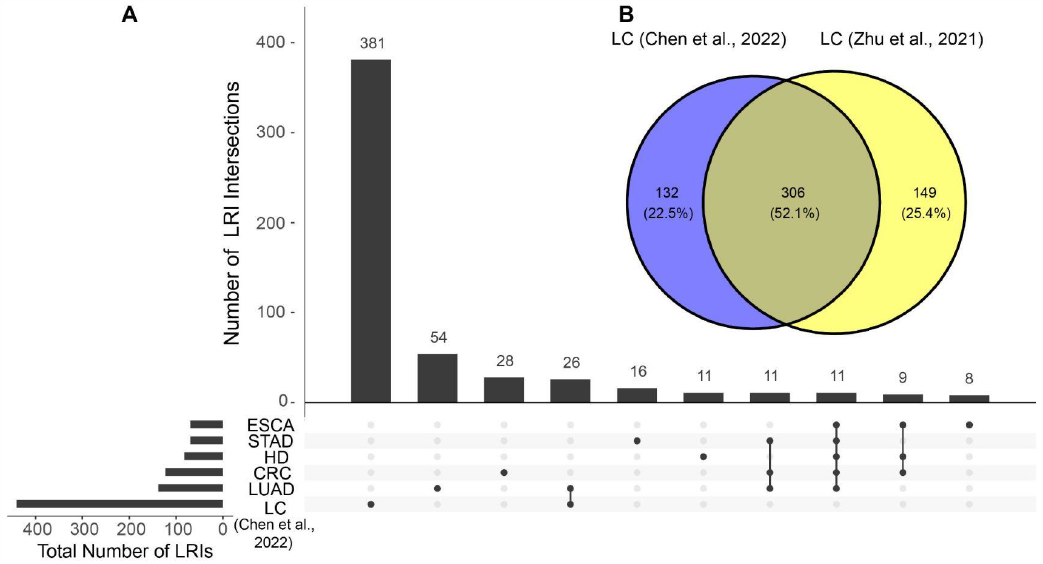
Similarities and differences of identified ligand-receptor interactions (LRIs) between cell-free RNA datasets. (**A**) LRI intersections in the cell-free RNA datasets from Chen *et al*.. Only the ten largest intersections are shown. (**B**) LRI intersection between cell-free liver cancer (LC) datasets of Chen *et al*. and Zhu *et al*.. HD, healthy donor; STAD, stomach adenocarcinoma; LUAD, lung adenocarcinoma; CRC, colorectal cancer; ESCA, esophageal cancer.

### Increase in the number of hepatocyte interactions in liver cancer blood samples

One of the main advantages of LRIs is the ability to infer cell-cell interactions and cellular networks. Considering that LRIs were associated with target and source cell types in the scRNA-seq data analysis results, we used the latter to infer changes in cell-cell interactions of cancer patients and healthy donors through blood samples. In order to accomplish this, we started out by analyzing LRIs in single-cell RNA-sequencing (scRNA-seq) data of liver cancer tissue samples (Suppl. Figure 1B; Suppl. File 1) (Lu et al., 2022). We found 1,120 LRIs in scRNA-seq data which were associated with 11 source and target cell types found in the liver (Suppl. File 1). Additionally, considering previous bulk RNA-sequencing findings which highlighted lowly expressed LRIs not captured by scRNA-seq data analysis (Villemin et al., 2023), we also identified LRIs in bulk RNA-sequencing data of liver cancer tissue samples to achieve a more comprehensive collection of LRIs. (Suppl. Figure 1B; Suppl. File 1) (Jiang et al., 2019; Zeng et al., 2020). Consistent with a prior observation (Villemin et al., 2023), a greater number of LRIs were detected in scRNA-seq data, with bulk RNA-sequencing contributing a modest number of unique LRIs (Suppl. Figure 1B).

We then visualized the top ten most abundant cell-cell interactions found in the scRNA-seq results for each cell-free RNA dataset. The analysis of the ten most abundant cell-cell interactions (Figure 2; Suppl. Figure 2; Suppl. File 2) showed strong hepatocyte signaling - both as a source and a target - in Chen *et al*. and Zhu *et al*. liver cancer datasets (Figure 2B; Suppl. Figure 2F). In contrast, hepatocyte signaling was less pronounced and ligands originating from hepatocytes were absent in healthy donor (Figure 2A; Suppl. Figure 2E) and other cancer blood samples (Suppl. Figure 2A-D). This aligns with prior studies describing an increase in hepatocyte signaling in the blood of liver cancer patients (Morlion et al., 2023; Vong et al., 2021).

**Figure 2:**
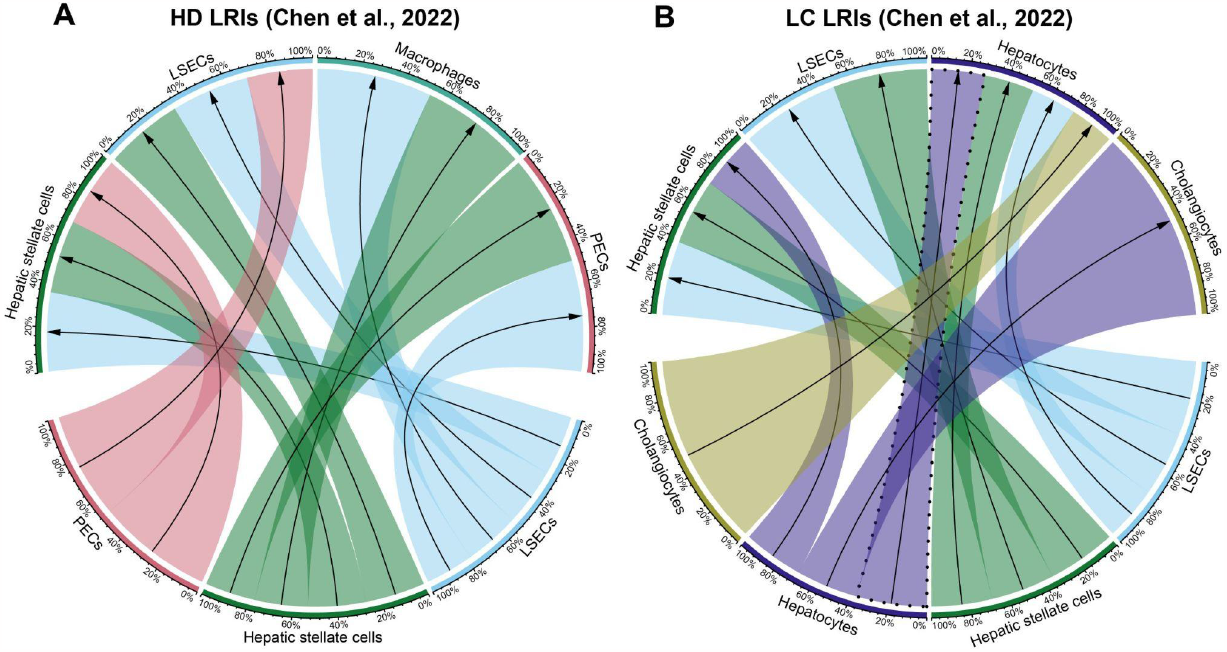
Cellular interactions in cell-free RNA datasets of Chen *et al*.. (**A**) Healthy donor (HD) and (**B**) liver cancer (LC) datasets. Visualized are the ten most abundant cell-cell interactions found in each dataset. Hepatocyte autocrine interactions are highlighted with a dashed line. Percentages represent the relative contribution of each interaction within the cumulative interactions for respective source and target cell types.

Another noteworthy observation was the detection of an abundant hepatocyte autocrine interaction in the liquid biopsy liver cancer in contrast to healthy control datasets. (Figure 2B; Suppl. Figure 2F). This finding confirms observations by previous studies indicating that the increase in autocrine hepatocyte signaling is a strong signal for liver carcinogenesis that can also be detected in the cell-free transcriptome (Ghoshdastider et al., 2021; Lu et al., 2022; Tummala et al., 2017).

### In-depth analysis of highly robust liver cancer LRIs

In order to more deeply understand the biological relevance of liver cancer-specific LRIs, our goal was to distill highly robust LRI signals. To achieve that, we created a panel of liver cancer-specific and reproducible LRIs, from the list of LRIs shared between Chen *et al*. and Zhu *et al*. cell-free liver cancer cohorts. In this panel, we excluded all LRIs found in other cell-free datasets to maximize the specificity towards liver cancer signals (Figure 3A). Next, to ensure that only LRIs remained which were previously reported to be present in liver cancer tissue, we filtered out the LRIs not found in either single-cell or bulk liver cancer RNA-seq datasets (Figure 3A). The final list consisted of 30 LRIs involving 22 ligands and 16 receptors (Figure 3B; Suppl. File 3). Notably, many of the identified ligands and receptors were previously associated with liver cancer. Among the candidates that were found, SERPINC1, GPC3 and the receptor ERBB3 were particularly noteworthy. SERPINC1 has a pronounced specificity to the liver and gallbladder (Xu et al., 2021) and its expression increases further in hepatocellular carcinoma (HCC) (Xu et al., 2021) and decreases during cholangiocarcinoma (ICC) (Suppl. Figure 3A) (Li et al., 2019; Wang et al., 2006) which also indicates its potential as a marker to differentiate liver cancer subtypes. GPC3 is an oncofetal marker for the liver whereas in the healthy adult liver little GPC3 expression has been detected (Morford et al., 2007; Zhou et al., 2018). Yet, upon cancerogenesis, the ligand is highly expressed in HCC (Suppl. Figure 3B) (Morford et al., 2007; Zhou et al., 2018). GPC3 promotes cancerogenesis via activation of the Wnt/β-catenin signaling pathway and, hence, presents itself as a potential therapeutic target (Kolluri and Ho, 2019; Shih et al., 2020). Finally, another noteworthy receptor from the list is ERBB3 (HERT3) which is upregulated in chronic hepatitis B (CHB) but not hepatitis C induced HCC (Buta et al., 2016) corresponding to the liquid biopsy liver cancer sample composition we employed (>60% CHB) (Chen et al., 2022).

**Figure 3:**
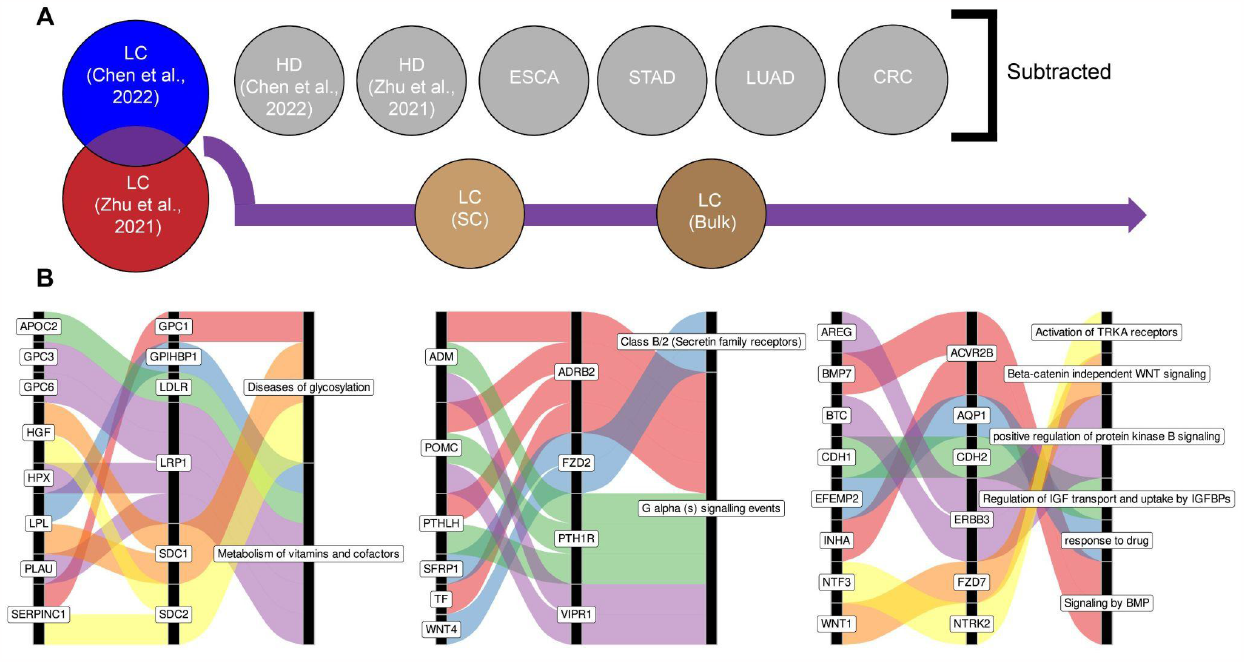
Curation of a panel of reproducible liver cancer specific ligand-receptor interactions (LRIs). (**A**) Schematic overview of the pipeline. Briefly, from the common LRIs between cell-free LC datasets of Chen *et al*. and Zhu *et al*. we excluded LRIs found in cell-free healthy donor (HD), esophageal cancer (ESCA), stomach adenocarcinoma (STAD), lung adenocarcinoma (LUAD) and colorectal cancer (CRC) datasets. Subsequently, LRIs not found in either liver cancer single-cell of bulk RNA-seq datasets were excluded as well. (**B**) Visualization of the identified panel of 30 LRIs and the largest regulated downstream pathway. For visualization purposes, the results are split into three alluvial plots.

Similarly, most of the downstream pathways regulated by the detected LRIs (Figure 3B) have previously documented relevance to liver pathologies (Ehata and Miyazono, 2022; Gao et al., 2021; Li et al., 2021; Mehta et al., 2015; Wang et al., 2021; Whittaker et al., 2010). The liver plays an important role in the metabolism of vitamins and various cofactors and dysregulation in these processes is commonly observed during liver pathologies (Fang et al., 2019; Licata et al., 2021; Paganoni et al., 2021; Raza et al., 2021; Roberts and Sarkar, 2008). Ectopic activation of WNT signaling is one of the main hallmarks of HCC (Khalaf et al., 2018; Kolluri and Ho, 2019; Perugorria et al., 2019; Wang et al., 2019) and protein kinase B (AKT) has been shown to be a key regulator of HCC progression (Khalaf et al., 2018; Nicholson and Anderson, 2002; Tian et al., 2023).

In addition, we endeavored to further contextualize the identified LRIs by using the CITE platform which assigns Relative Crosstalk (RC) scores to LRIs based on the expression patterns of the LRIs in the tumor microenvironment (Ghoshdastider et al., 2021). Of the 30 LRIs present in the database, 11 had been assigned RC scores by CITE in HCC (Suppl. Figure 4), with the vast majority having low RC scores in the normal cell compartment which suggests a higher association of these LRIs with cancer. The majority of the LRIs also had a high RC score in the stromal compartment, which includes autocrine LRIs between stromal cells and paracrine LRIs between stromal and cancer compartments, in line with previous findings (Ghoshdastider et al., 2021). Here, the INHA-ACVR2B LRI stands out given its high cancer compartment autocrine RC score which is in line with a prior observation of ACVR2B interactions being very prevalent in cancer-cancer communication across different tumor types (Ghoshdastider et al., 2021) and lends further support to our observation of a potential hepatocyte autocrine signaling. Together, these findings suggest that the cell-free RNA signals in the blood of liver cancer patients can be utilized to indirectly gain insights into changes in ligand-receptor interactions during liver carcinogenesis.

## Data availability

Publicly available datasets analyzed during the study are available from the NCBI GEO repository under the accession numbers GSE174302, GSE142987, GSE149614 and GSE124535.

## Code availability

The computational code used in the study is available at GitHub: https://github.com/aramsafrast/cfLRIs/

## Author contributions

AS and DW conceptualized the study. AS performed all data analyses. AS and DW wrote the manuscript.

