## Supplementary Figures for "Detection of reproducible liver cancer specific ligand-receptor signaling in blood"

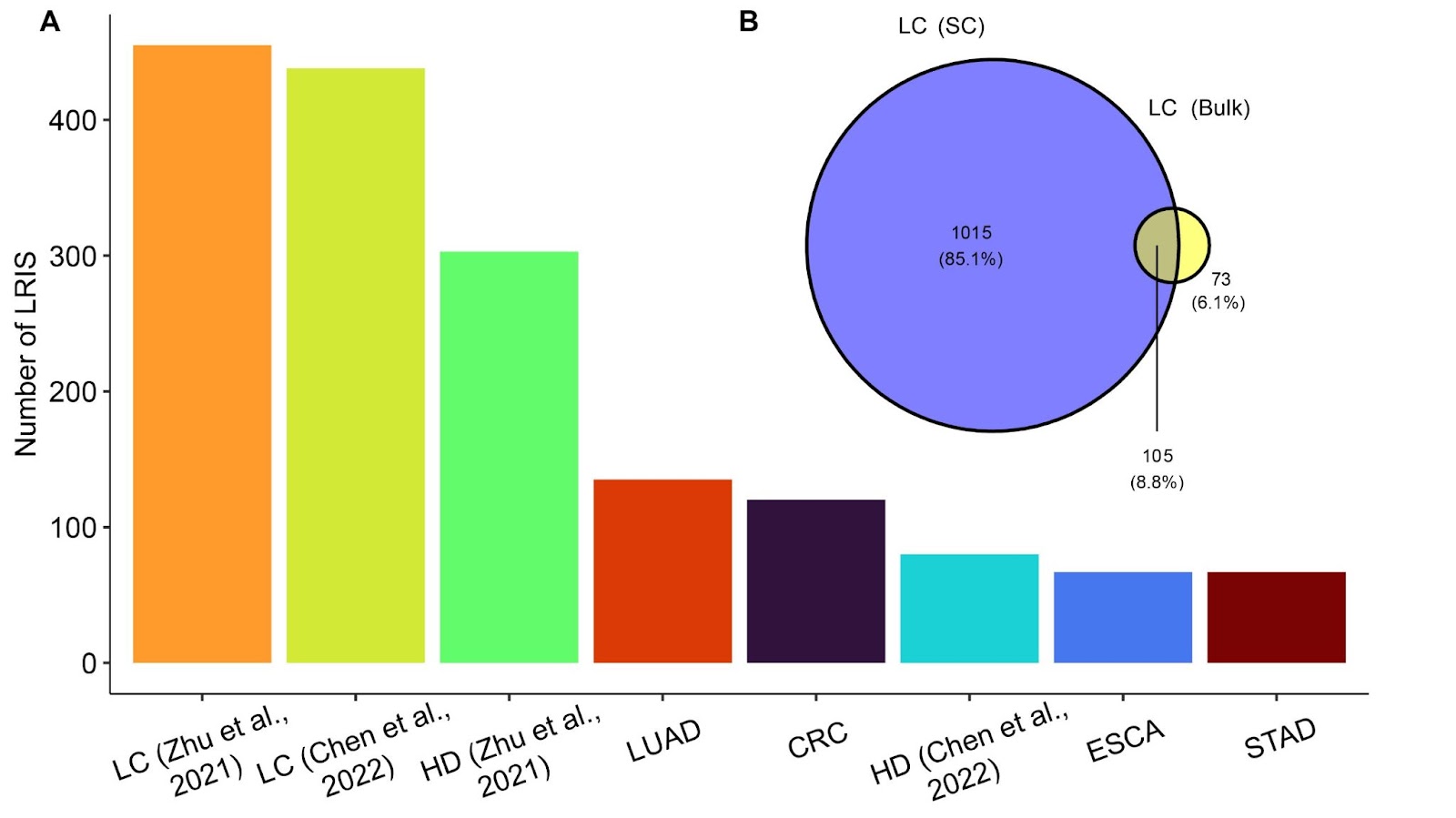
 **Suppl. Figure 1**: **Total number of ligand-receptor interactions (LRIs) found across datasets**. (**A**) Number of LRIs in cell-free RNA datasets of Chen *et al*. and Zhu *et al.*. (**B**) The number of unique and common LRIs between liver cancer (LC) tissue single-cell (SC) and bulk RNA-seq datasets. HD, healthy donor; LUAD, lung adenocarcinoma; CRC, colorectal cancer; ESCA, esophageal cancer; STAD, stomach adenocarcinoma.

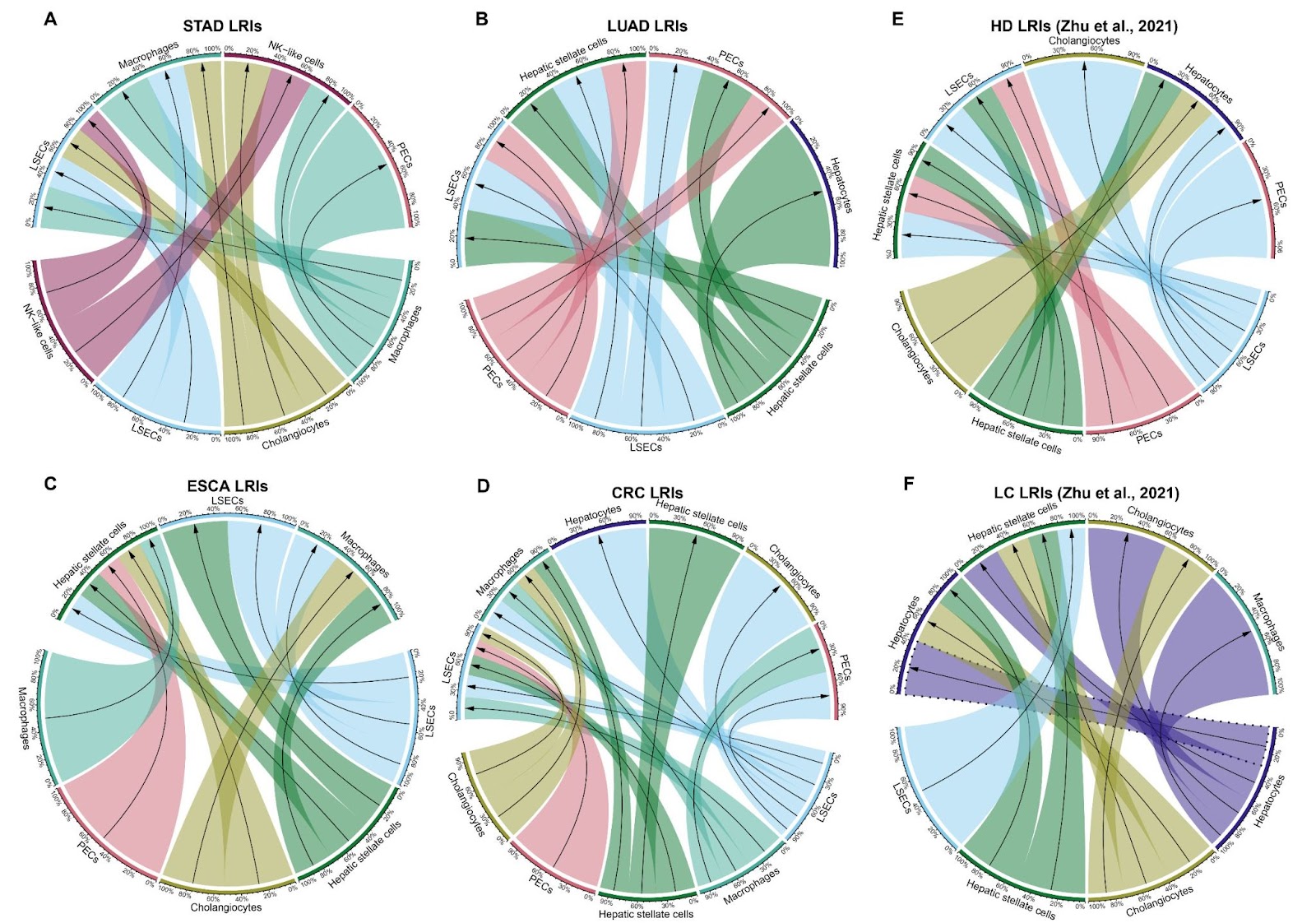

**Suppl. Figure 2**: **Cellular interactions in cell-free RNA datasets of Chen *et al***. **and Zhu *et al***.. (**A**) Stomach adenocarcinoma (STAD), (**B**) Lung adenocarcinoma (LUAD), (**C**) esophageal cancer (ESCA), (**D**) colorectal cancer (CRC) datasets from Chen *et al*., (**E**) healthy donor (HD) and (**F**) liver cancer (LC) datasets from Zhu *et al*.. Visualized are the ten most abundant cell-cell interactions found in each dataset. Hepatocyte autocrine interactions are highlighted with a dashed line. Percentages represent the relative contribution of each interaction within the cumulative interactions for respective source and target cell types.

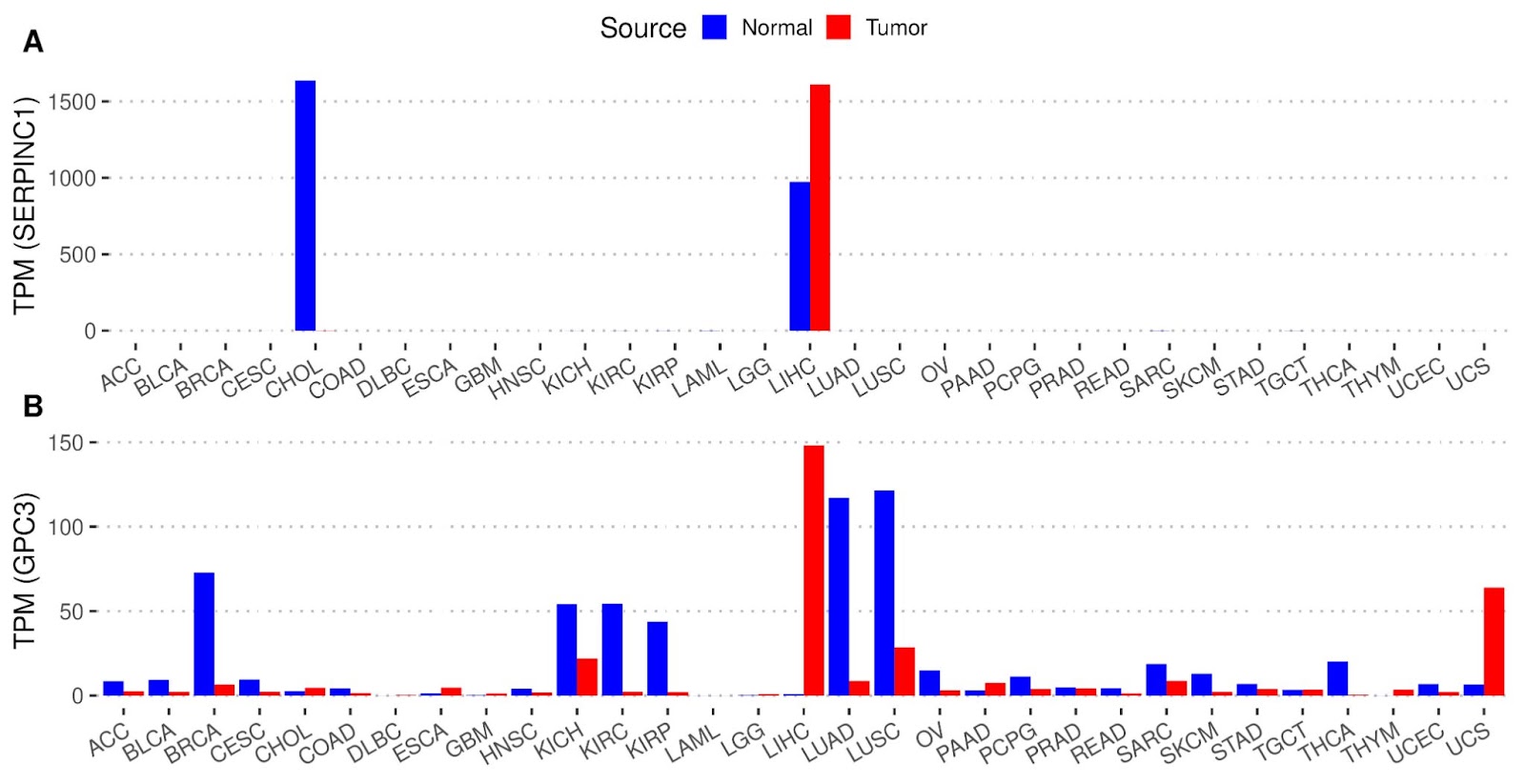

**Suppl. Figure 3**: **Gene expression of ligands (A) SERPINC1 and (B) GPC3 across paired normal and tumor tissues**. Data was collected from the online portal GEPIA2. ACC, Adrenocortical carcinoma; BLCA, Bladder Urothelial Carcinoma; BRCA, Breast invasive carcinoma; CESC, Cervical squamous cell carcinoma and endocervical adenocarcinoma; CHOL, Cholangiocarcinoma; COAD, Colon adenocarcinoma; DLBC, Lymphoid Neoplasm Diffuse Large B-cell Lymphoma; ESCA, Esophageal carcinoma; GBM, Glioblastoma multiforme; HNSC, Head and Neck squamous cell carcinoma; KICH, Kidney Chromophobe; KIRC, Kidney renal clear cell carcinoma; KIRP, Kidney renal papillary cell carcinoma; LAML, Acute Myeloid Leukemia; LGG, Brain Lower Grade Glioma; LIHC, Liver hepatocellular carcinoma; LUAD, Lung adenocarcinoma; LUSC, Lung squamous cell carcinoma; OV, Ovarian serous cystadenocarcinoma; PAAD, Pancreatic adenocarcinoma; PCPG, Pheochromocytoma and Paraganglioma; PRAD, Prostate adenocarcinoma; READ, Rectal adenocarcinoma; SARC Sarcoma; SKCM, Skin Cutaneous Melanoma; STAD, Stomach adenocarcinoma; TGCT, Testicular Germ Cell Tumors; THCA, Thyroid carcinoma; THYM, Thymoma; UCEC, Uterine Corpus Endometrial Carcinoma; UCS, Uterine Carcinosarcoma; TPM, transcripts per million.

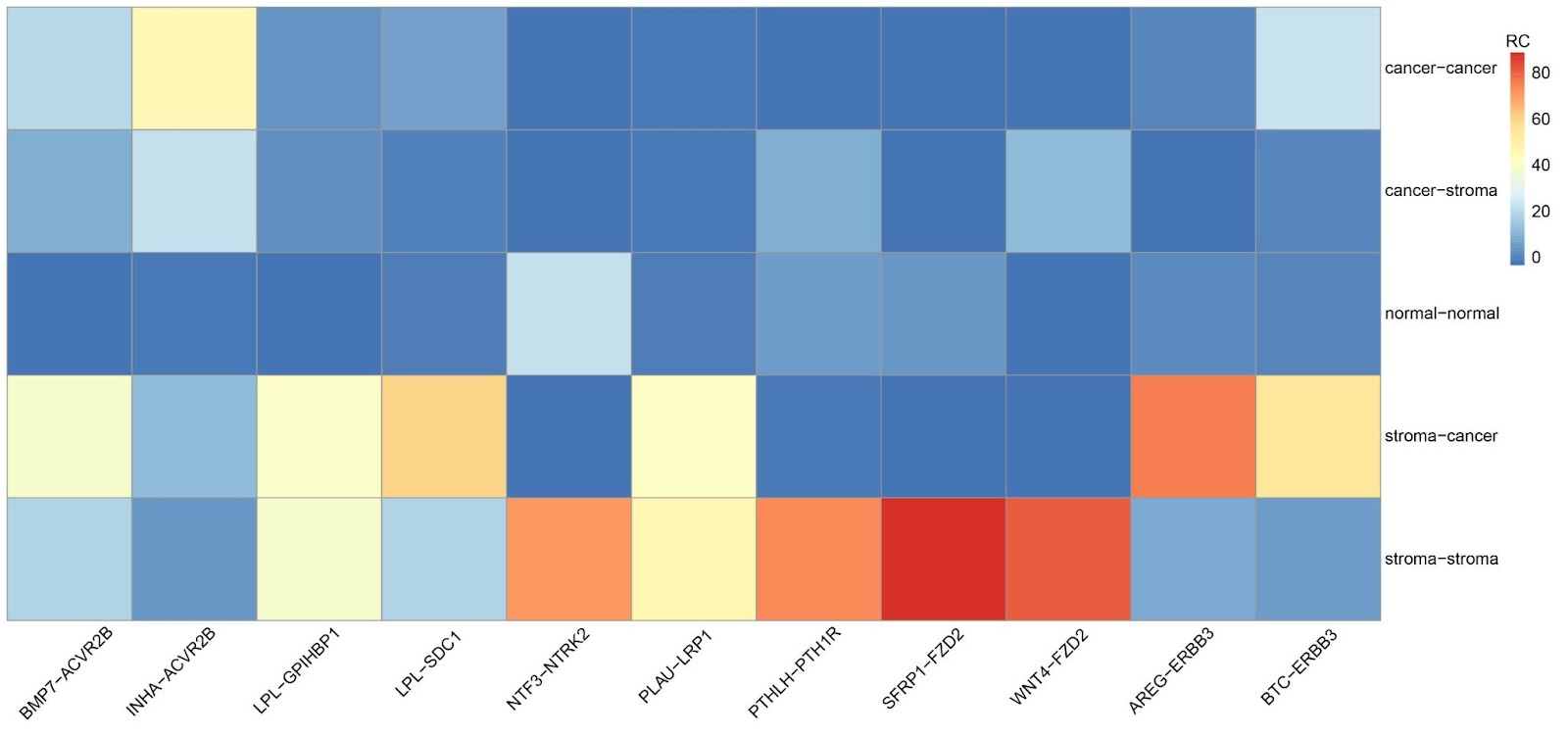

**Suppl. Figure** **4**: **Relative Crosstalk (RC) scores of 11 ligand-receptor interactions (LRIs) across tumor microenvironment compartments in liver cancer**. RC scores measure the relative autocrine and paracrine crosstalk between cancer and stromal cell compartments in the tumor microenvironment.
